# Investigation of extramammary sources of Group B *Streptococcus* reveals its unusual ecology and epidemiology in camels

**DOI:** 10.1101/2021.05.27.445946

**Authors:** Dinah Seligsohn, Chiara Crestani, Nduhiu Gitahi, Emelie Lejon Flodin, Erika Chenais, Ruth N. Zadoks

## Abstract

Camels are vital to food production in the drylands of the Horn of Africa, with milk as their main contribution to food security. A major constraint to camel milk production is mastitis, inflammation of the mammary gland. The condition negatively impacts milk yield and quality as well as household income. The leading cause of mastitis in dairy camels is *Streptococcus agalactiae,* group B *Streptococcus* (GBS), which is also a commensal and pathogen of humans. It has been suggested that extramammary reservoirs for this pathogen may contribute to the occurrence of mastitis in camels. We explored the molecular epidemiology of GBS in camels using a cross-sectional study design for sample collection and phenotypic, genomic and phylogenetic analysis of isolates. Among 88 adult camels and 93 calves from six herds in Laikipia County, Kenya, GBS was detected in 20% of 50 milk samples, 25% of 152 nasal swabs, 8% of 90 oral swabs and 3% of 90 rectal swabs, but not in vaginal swabs. Per camel herd, two to four sequence types (ST) were present. More than half of the isolates belonged to ST617 or its single-locus variant, ST1652, with these STs found across all sample types. Serotype VI was detected in 30 of 58 isolates. In three herds, identical STs were detected in milk and swab samples, suggesting that extramammary sources of GBS may contribute to the maintenance and spread of GBS within camel herds. This needs to be considered when developing prevention and control strategies. In addition, the high nasal carriage rate, low recto-vaginal carriage rate, and high prevalence of serotype VI for GBS in camels are in stark contrast to the distribution of GBS in humans and reveal hitherto unknown ecological and molecular features of this bacterial species.

## Introduction

In the arid and semi-arid lands of the Horn of Africa, nomadic pastoralism is common, and livestock is mainly kept for sustenance [1]. Camels are well-adapted to surviving despite limited access to water and feed [2] and have long been kept for milk and meat by pastoralist communities in this region [1]. The importance of camels is likely to grow because of climate change, resulting in prolonged and recurrent droughts and erratic rainfall [3]. Because of this, camel keeping is increasing in traditional cattle-keeping communities in Kenya [4]. Furthermore, the demand for camel milk has amplified over the past decades. Consequently, the camel milk sector in Kenya is currently undergoing substantial changes despite being hampered by lack of resources and adequate infrastructure [5].

Mastitis, inflammation of the udder most often caused by bacterial infection, is a common constraint to milk production in dairy camel herds in this region [6,7]. This has implications for household income and food security, as well as animal and public health [8]. *Streptococcus agalactiae*, also referred to as Group B *Streptococcus* (GBS), is one of the most important udder pathogens in camels, often resulting in chronic infections [7]. In dairy cattle, GBS was considered an obligate intramammary pathogen, but this paradigm has recently been challenged as GBS has been found in the environment and body sites other than the mammary gland (e.g. in faeces and in swabs taken from the rectum, vagina and throat) of cattle in South America, North America and Europe [9–11]. Similarly, in camels, GBS has been isolated from the nasal and vaginal mucosa of clinically healthy camels [12], exemplifying extramammary sources of GBS that may constitute a reservoir for intramammary infections. In cattle, cross-suckling by calves may lead to pathogen transmission from milk via the oral mucosa of one calf to the juvenile udder of another calf, causing intramammary GBS infection [13]. Cross-suckling behaviour by calves is seen in housed animals, a management system that is not used in East African camels. In this region, camel calves, are kept together with their mothers on pasture [14], and cross-suckling of dams by calves could contribute to the spread of GBS if they carry the organism on mucosal surfaces. Although Fischer et al. [15] found that camel GBS isolates were associated with various clinical presentations and body sites, the role of healthy carriers in maintaining a GBS population, and their involvement in GBS mastitis epidemiology have not been investigated.

In this study we used phenotypic, genomic and phylogenetic methods to investigate GBS diversity in isolates from camel milk and extramammary sources, with a specific focus on the potential role of extramammary strains in mastitis and the overall objective to expand the knowledge basis for mastitis control in camels.

## Materials and methods

### Ethics statement

The study was approved by the National Commission for Science, Technology and Innovation, Nairobi, Kenya (Permit number: NACOSTI/P/19/84995/13088). Prior to sampling, animal owners and herders received oral or written information about the purpose of the study and the sampling procedures, and consent to participate was obtained. Participants were informed that inclusion in the study would be anonymous and that they could withdraw from the study at any time. No additional permissions for sample collection and analysis were required.

### Study area and selection of herds

The study was conducted in Laikipia County, Kenya. Laikipia is classified as semi-arid with an average annual rainfall of 639 mm. The land is predominantly used for ranching and other forms of animal production in the north, including large-scale ranches with hired labour and subsistence smallholder farmers using communal land for grazing, with permanent subsistence agriculture dominating in the south [16]. The total camel population measures 9,800 [17] and acquisition of camels among pastoralist groups without a tradition of camel keeping is increasing [18]. Most camels are semi-sedentary. The study area was selected because of its considerable and growing camel population, and the presence of a camel milk collection center. Sampling took place during the wet season, November 2019. Herds (n=6, A-F) were selected on the basis of prior knowledge of prevalent subclinical mastitis [19].

### Sampling and bacteriological culture

Sampling was carried out in connection with the early morning milking between 4 am and 8 am. Calves were sampled before being released to suckle their mother to initiate milk let down. All lactating camels were screened using the California Mastitis Test [20], where viscosity is an indicator of the severity of inflammation and scored according to the Nordic scale (1-5) [21], where 1 signifies no change in viscosity and 5 signifies a marked increase in viscosity with formation of a distinct gel peak. From camels with a CMT-score of 2 or higher, composite milk samples were collected aseptically in sterile plastic 50 mL vials. Milk samples were kept cold during transport using cool bags and frozen at −20°C within 4 hours from the time of collection. Swab samples were collected from all lactating camels and their calves using sterile flocked nylon swabs (e-swab, COPAN diagnostics Inc. Murrieta, CA, US). Prior to swabbing, a clinical assessment of each sampling site was carried out. Clinical abnormalities, such as nasal or vaginal discharge, swelling, lesions or diarrhoea were noted. Swabs were taken from the nasal and vaginal mucosa of lactating camels, and from the nasal, oral and rectal mucosa of their respective suckling calves. The nasal mucosa was sampled by flaring the nostrils open and swabbing the inside of the nasal cavity of both nostrils with the same swab. The vaginal mucosa was sampled by separating the labia, inserting the swab and rolling it against the vaginal wall. Oral samples were collected by inserting the swab into the oral cavity of the calf and rolling it against the buccal and pharyngeal mucosa. The rectal mucosa was sampled by inserting the swab in the rectum and gently rolling it against the rectal wall. Swab samples were kept refrigerated at 4-8°C and processed at the Department of Public Health, Pharmacology and Toxicology, University of Nairobi, Nairobi, Kenya, within 1 to 10 days from collection.

For processing, milk samples were thawed at room temperature. Ten μL of milk were cultured on modified Edward’s agar (EA) (Oxoid, CM0027), and incubated aerobically for 24 and 48 hours at 37°C before final examination. Milk samples that were negative for GBS on direct culture were enriched in Todd Hewitt (TH) broth (Oxoid, CM0189) at 37°C for 18 to 24 hours and re-cultured on EA. Catalase-negative, KOH-negative colonies, displaying blue pigmentation, with or without β-haemolysis, were subcultured on blood agar (Oxoid, CM0271) containing 5% defibrinated sheep blood.

Swab samples were enriched in TH broth at 37°C for 18-24 hours prior to plating. A calibrated loop (10 μL) was used to plate the enrichment on EA and the plates were incubated aerobically at 37°C for 18-48 hours. Colonies that showed the same phenotypic and biochemical characteristics as described above were subcultured on EA and incubated at 37°C for 18-48 hours. All CAMP (Christie-Atkins-Munch-Peterson)-positive [22] bacterial isolates (n=58) were confirmed as GBS using a slide latex agglutination test (Streptex Latex Agglutination Test, ThermoFisher Scientific Inc., Waltham, MA, USA). GBS isolates were subcultured on blood agar to assess purity and inoculated on stab agars (SVA, Uppsala, Sweden) that were incubated aerobically for 8-12 hours in 37°C before storage at 4-8°C. Stab agars were transported at ambient temperature to the National Veterinary Institute (SVA), Uppsala, Sweden, which took 24 hours, and then recultured on blood agar containing 5% defibrinated horse blood (SVA, Uppsala, Sweden) and incubated aerobically for 24-48 hours at 37°C. Species identity was confirmed using matrix assisted laser desorption ionization-time of flight mass spectrometry analysis (MALDI-ToF MS) [23]. The bacteria were analyzed in duplicates. Criteria for species identification were as follows: a score of ≥2 indicated identification at species level, 1.80 to 1.99 at genus level, and <1.80 no identification. Species identification was performed using a custom-made database including the Bruker databases no. 5627 and no. 5989.

### DNA extraction and sequencing

Extraction of DNA was carried out from all confirmed GBS isolates (n=58), using a magnetic bead-based method. A calibrated loop (1 μL) was used to suspend colony material in 600 μL nuclease free water (Sigma-Aldrich, St Louis, MO, USA) and mixed with 0.1 mm silica beads (BioSpec Products Inc., Bartlesville, USA). The suspension was added to the FastPrep24 (MP Biomedicals LLC, Irvine, CA, USA) and run at 6.5 m/s for three 2-minute cycles. DNA was extracted from 200 μL samples using the IndiMag Pathogen kit (Indical Bioscience GmbH, Leipzig, Germany) and eluted in nuclease free water. The Invitrogen Qubit 3.0 Fluorometer and the Qubit dsDNA BR Protein Assay kit (ThermoFisher Scientific Inc., Waltham, MA, USA) were used to measure DNA concentrations, which were adjusted to 7.5 ng/μL. Library preparation and whole genome sequencing were performed by Clinical Genomics, Science for Life Laboratory (Clinical Genomics, Solna, Sweden) on the Illumina NovaSeq (Illumina, Inc. CA, US), resulting in paired-end libraries of 150bp read length.

### Antimicrobial susceptibility testing

GBS isolates were tested for presence of phenotypic antimicrobial resistance by determination of minimum inhibitory concentration (MIC). Testing (but not interpretation) was performed according to standards of the Clinical and Laboratory Standards Institute [24] using a broth microdilution method, cation-adjusted Mueller-Hinton broth, Sensititre™ STAFSTR panels (TREK Diagnostic System, UK) and Sensititre™ NLD1GNS panels (TREK Diagnostic System, UK). As quality control, a strain of *Staphylococcus aureus* ATCC 15019 was tested in parallel with the isolates; results were within acceptable ranges. The MIC were determined for the following compounds: cephalotin, clindamycin, enrofloxacin, erytromycin, gentamicin, nitrofurantoin, penicillin, tetracycline and trimethoprim-sulfametoxazol. GBS specific epidemiological cut-off (ECOFF) values issued by the European Committee on Antimicrobial Susceptibility Testing (EUCAST; clindamycin, erythromycin, nitrofurantoin, penicillin and tetracycline) were used to classify isolates as wild type (WT; the ECOFF based counterpart of susceptibility, which is defined based on clinical cut-off values) or non-wild type (non-WT, the ECOFF-based counterpart of resistance), with clinical breakpoints from SVA used for the remaining compounds (cephalotin and trimethoprim-sulfametoxazol). *Streptococcus* species have a low inherent susceptibility to quinolones and an ECOFF value is not defined by EUCAST. MIC for enrofloxacin was determined in this study but not classified as WT, non-WT, susceptible or resistant due to the lack of available cut-off values. Seventeen isolates showed growth at the highest concentration of gentamicin provided at 8 μg/mL and were subsequently retested on the Sensititre™ NLD1GNS panel with a wider range for gentamicin (up to 32 μg/mL). SRST2 was used to detect antimicrobial resistance (AMR) genes from raw reads with the ARG-ANNOT v3 database [25].

### Lactose typing

Lactose fermentation was assessed phenotypically by inoculating each of the selected isolates onto bromocresol purple lactose agar (SVA, Uppsala, Sweden). Lactose fermentation was indicated by a yellow colour change, whereas the absence of colour change was interpreted as negative for lactose fermentation. *Escherichia coli* ATCC35218 and *Proteus mirabilis* CCUG26767 were used as positive and negative controls, respectively. Plates were incubated aerobically at 37°C and checked for colour change at 24, 48, and 72 hours and after one week of incubation. The presence of the lactose operon (Lac.2) [26] was investigated with a BLASTn v2.9.0 search [27] based on a database of five known Lac.2 genotypic variants [28,29]. Thresholds for positivity were set at 90% for both sequence identity (ID) and query coverage (QC). Lac.2-negative genomes were confirmed by manual scanning of annotation files obtained using Prokka v1.14.6 [30], searching for genes associated with the metabolism of lactose. To further support these findings, we conducted a PCR targeting a ≈2.5-kbp region straddling *lacEG*. Positive and negative controls were selected from among the study isolates based on genomic detection of Lac.2.

### Phylogenetic and statistical analysis

Reads were filtered and trimmed with ConDeTri suite v2.3 [31]. *De novo* assembly was performed using SPAdes v3.13.1 [32] and assembly quality was checked with QUAST v5.0.2 [33]. Bacterial species identity was confirmed with KmerFinder v3.2 [34]. All assembled genomes passed quality control. Multi locus sequence typing (MLST) was carried out with SRST2 v0.2.0 [35] and capsular serotype was detected *in silico* using a standard method [36].

Snippy v4.6.0 (https://github.com/tseemann/snippy) was used for alignment of the core genome. ILRI112 (accession HF952106), which is an ST617 genome from a camel from Kenya, was used as reference. A maximum likelihood tree was inferred using RAxML-NG v0.9.0 [37] under a GTR+G model. A map of herd coordinates was created with ggmap with R, in RStudio (v4.0). All figures were edited using Inkscape (www.inkscape.org).

Data editing was carried out in Excel (Microsoft Corp., Redmond, WA). Associations between categorical variables were tested for statistical significance using Pearson’s chi-square test or Fisher’s exact test when suitable. A *p-*value of 0.05 or lower was considered significant. All statistical analyses were performed in STATA (Stata Statistical Software, release 13.1; StataCorp LP, College Station, TX).

## Results

In all, 50 milk samples, 88 nasal mucosa swabs and 88 vaginal mucosa swabs were collected from 88 adult camels in six herds (A to F). Swab samples (n=244) were also collected from the nasal (n=64), oral (n=90) and rectal (n=90) mucosa of 93 suckling calves. Due to occasional difficulties in restricting camels for sampling or camels being released on pasture before the sampling was completed, some sample categories are missing for a few individuals in herds A, D, E and F.

### Herd demographics

Herds belonged to different management systems, but all camels were browsed during the day and kept overnight in “bomas” (an animal enclosure traditionally made from twigs and branches or other more solid materials (**Table 1**)) with lactating dams separated from their calves. Three of the herds (A, B and D) were kept under ranch management, i.e. herds belonged to private landowners with no tradition of keeping camels who employed workers to handle their livestock. One herd (C) was semi-commercial, owned by pastoralists, kept under traditional pastoralist conditions but with employed herders. Two smaller herds (E-F) were managed by families and kept at the homestead, together with other types of animals. Milk from the smallholder herds was used for household consumption. Management characteristics are shown in **Table 1.** Estimated average herd size was 87 and the average number of lactating camels seventeen. Camels were mainly of Somali breed (n=62) or Turkana breed (n=15), followed by crossbreeds (n=6) and other breeds (n=4), information on breed was missing for one camel. Herd location is shown in **Figure 1**.

**Table 1.** Herd demographics and management characteristics of six dairy camel herds from a study into carriage and shedding of group B *Streptococcus* conducted in November 2019 in Laikipia County, Kenya.

| Herd and<br>management<br>characteristics | Herd |  |  |  |  |  |
| --- | --- | --- | --- | --- | --- | --- |
|  | A | B | C | D | E | F |
| **Management system** | Ranch | Ranch | Pastoralist | Ranch | Smallholder | Smallholder |
| **Mobile/permanent boma** | Mobile | Mobile | Mobile | Permanent | Permanent | Permanent |
| **Boma material** | Metal fence | Acacia | Acacia | Wooden fence | Acacia | Acacia |
| **Total herd size<sup>1</sup>** | 76 | 75 | 180 | 86 | 53 | 53 |
| **Number of lactating camels** | 19 | 23 | 18 | 16 | 17 | 8 |
| **Milking frequency (times/day)** | 1 | 2 | 2 | 4 | 2 | 2 |
| **Post-milking cleaning of hands<sup>2</sup>** | Yes | Yes | No | No | No | No |
| **Milking order** | No | No | Newly calved camels milked first | Oldest camels milked first | Newly calved camels milked first | Newly calved camels milked first |
| **Selling milk** | Yes | Yes | Yes | Yes | Yes | No |
| **Watering interval (days)** | 2 | 0 | 1 | 0 | 1 | 0 |
| **Other animals kept in contact with camel herd** | Cattle, poultry, sheep | No | No | No | Cattle, poultry, sheep, goats, dogs | Cats, poultry, sheep, goats, dogs |
| **New camels added to the herd within the last year** | No | No | No | Yes | No | No |

<sup>1</sup>For herds not belonging to ranches, herd size was regarded as confidential, and a rough estimate was given by the herders.
<sup>2</sup>No pre-milking washing of hands was carried out in any of the herds.

**Figure 1.**
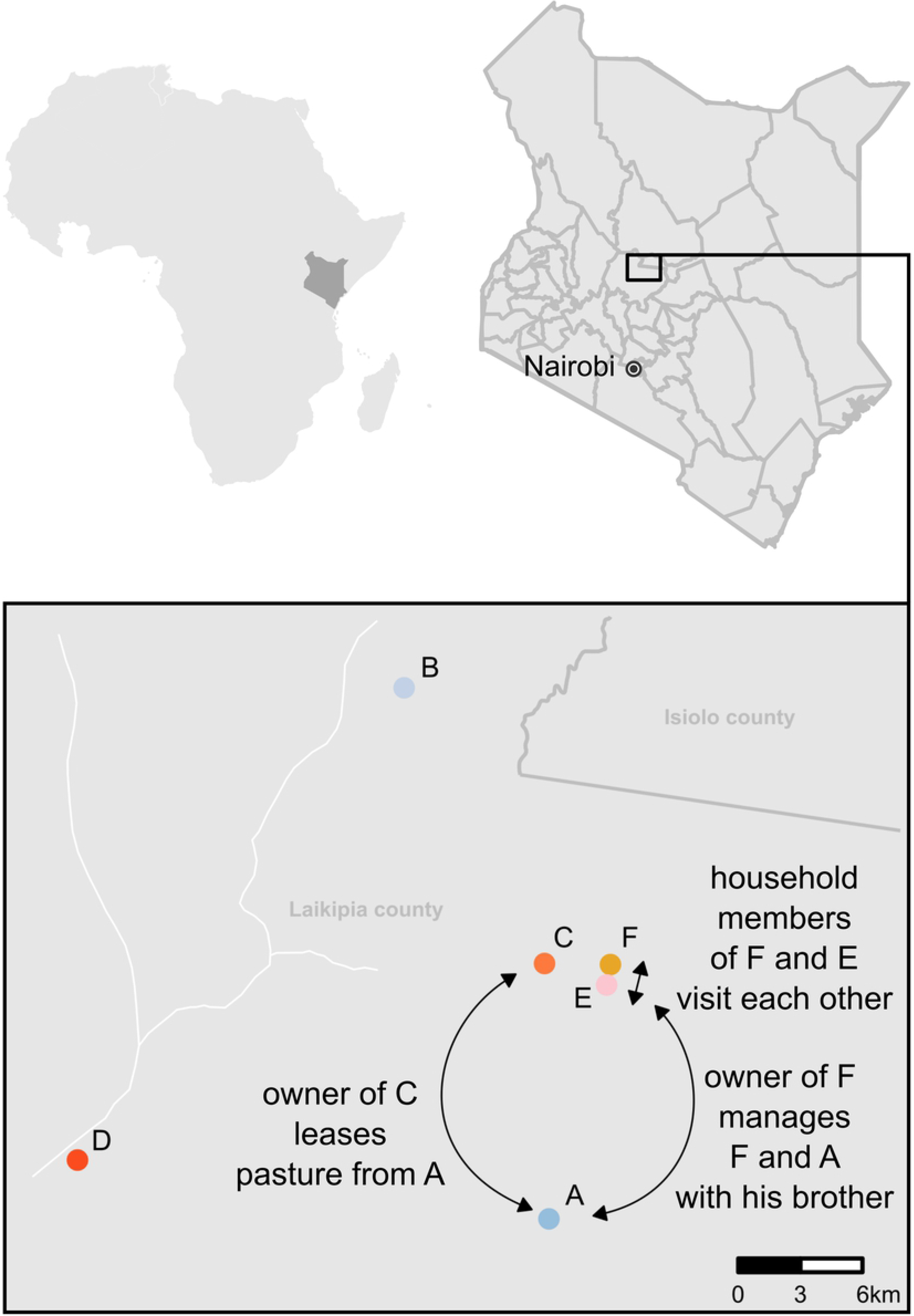
Geographical location of six dairy camel herds (A-F) in Laikipia County, Kenya. Arrows indicate known epidemiological links between herds. The map of herd coordinates was created with ggmap with RStudio in R (v4.0).

### Mastitis and GBS

Milk samples originated from animals with CMT ≥2 in at least one quarter, which was observed in 50 of 88 camels (57%), and in each herd. Most of these mastitis cases were subclinical, only nine camels displayed symptoms of acute mastitis, such as swollen udder, pain or deviations of the milk appearance (changes in colour, consistency, clotting or blood). Previous udder problems were reported for 25 of 88 camels and ten of 88 camels had at least one blind (non-milk producing) quarter. Four calves from three herds were observed to suckle from camels other than their mother (cross-suckling). GBS was isolated from milk of ten (20%) CMT-positive camels in four herds; seven of these camels had reportedly had long lasting udder problems, and there was a positive association between a previous history of mastitis and isolation of GBS from milk (*p*=0.004).

### Extramammary sources of GBS

Extramammary GBS was detected in all six herds, in 24 of 88 nasal swabs from adult camels and in 19 of 95 calves, including 14, 7 and 3 nasal, oropharyngeal and rectal swabs, respectively (**Table 2**). All swabs were taken from clinically healthy sampling sites with the exception of the presence of orf-like lesions on the muzzle of all calves in herd C. The most common site of GBS isolation was the nasal mucosa (38/151 samples). Only seven oral swabs from three herds and three rectal swabs from two herds were positive for GBS. No vaginal swabs were positive for GBS. In two herds (E and F), GBS was exclusively detected in extramammary samples.

**Table 2.**
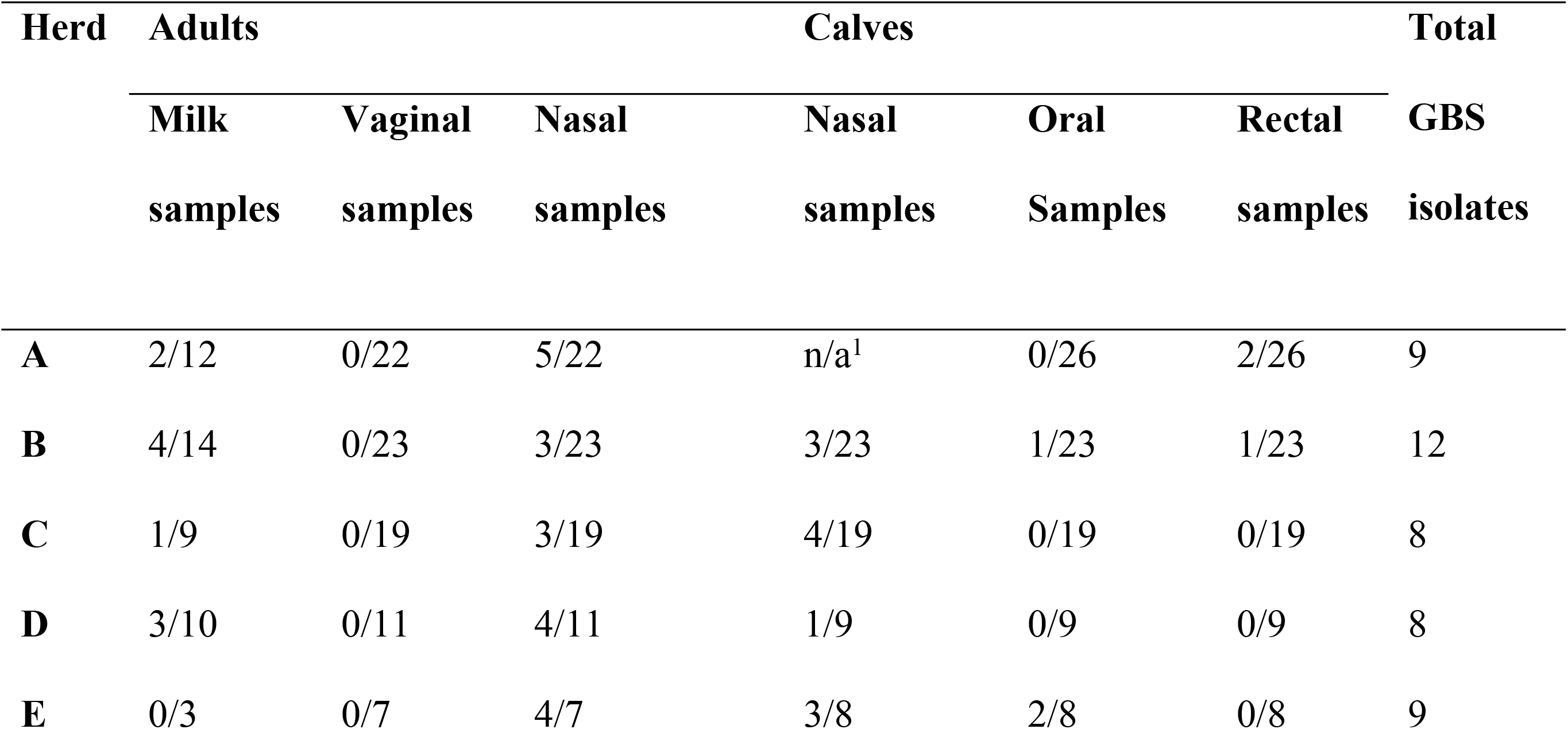
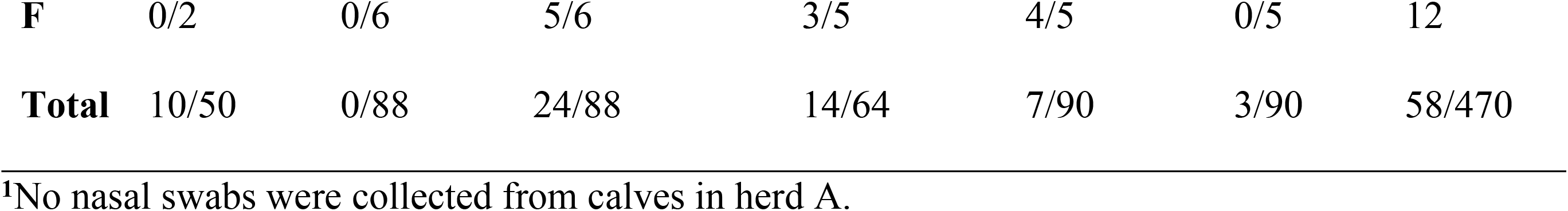
Group B *Streptococcus* (GBS) detected in milk and extramammary body sites of camels from six herds in Laikipia, Kenya, November 2019. (GBS-positive samples/total number of samples).

| Herd | Adults |  |  | Calves |  |  | Total |
| --- | --- | --- | --- | --- | --- | --- | --- |
|  | Milk samples | Vaginal samples | Nasal samples | Nasal samples | Oral Samples | Rectal samples | GBS isolates |
| A | 2/12 | 0/22 | 5/22 | n/a<sup>1</sup> | 0/26 | 2/26 | 9 |
| B | 4/14 | 0/23 | 3/23 | 3/23 | 1/23 | 1/23 | 12 |
| C | 1/9 | 0/19 | 3/19 | 4/19 | 0/19 | 0/19 | 8 |
| D | 3/10 | 0/11 | 4/11 | 1/9 | 0/9 | 0/9 | 8 |
| E | 0/3 | 0/7 | 4/7 | 3/8 | 2/8 | 0/8 | 9 |
| **F** | 0/2 | 0/6 | 5/6 | 3/5 | 4/5 | 0/5 | 12 |
| **Total** | 10/50 | 0/88 | 24/88 | 14/64 | 7/90 | 3/90 | 58/470 |

<sup>1</sup>No nasal swabs were collected from calves in herd A.

For nine camels, GBS was isolated from two sampling sites in the same individual (adults: n=3; calves: n=6).

### Diversity of sequence types and serotypes

One DNA extract did not pass quality control for sequencing and was excluded from analysis. Five sequence types (ST) were identified, all belonging to previously described camel-associated clonal complexes. Almost half of the isolates belonged to ST617 (n=17) or its single locus variant (SLV) ST1652 (n=19), followed by ST615 (n=10), ST612 (n=6) and ST616 (n=5). Within-herd diversity varied between herds: in one herd (A), isolates belonged to four STs, in two herds (D and F), isolates belonged to three STs and in the remaining three herds (B, D and E), isolates belonged to two STs (**Figure 2**). Within-host diversity was found in six camels, in which isolates belonged to two STs. There was a significant association between ST and sampling site (*p*=0.005). For example, four of five ST616 isolates were found in milk, and five of six ST612 isolates and 15 of 17 ST617 isolates were found in nasal swabs (**Table 3)**. ST1652 was isolated from all GBS-positive sample types whereas ST617 was found in nasal and oral swabs and in milk, ST612 in nasal and oral samples, and ST616 milk samples and an oral sample (**Figure 2**.).

**Table 3.**
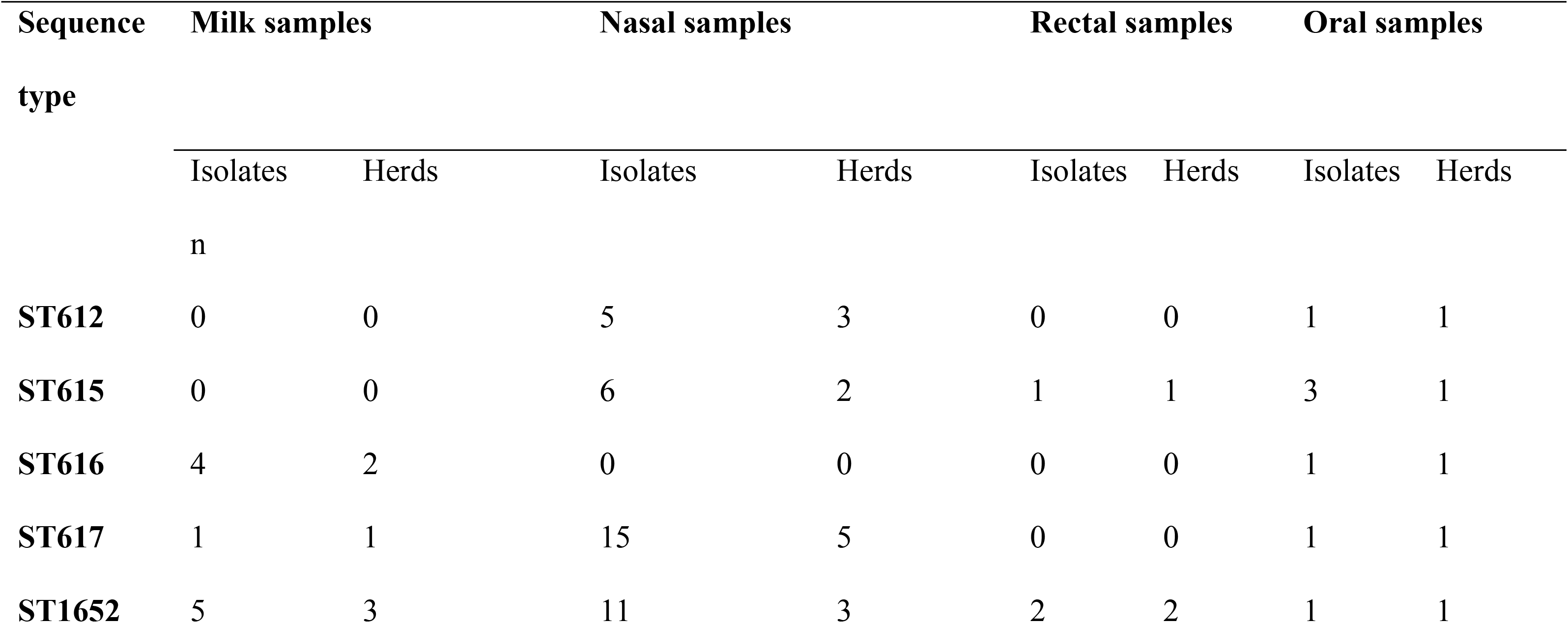
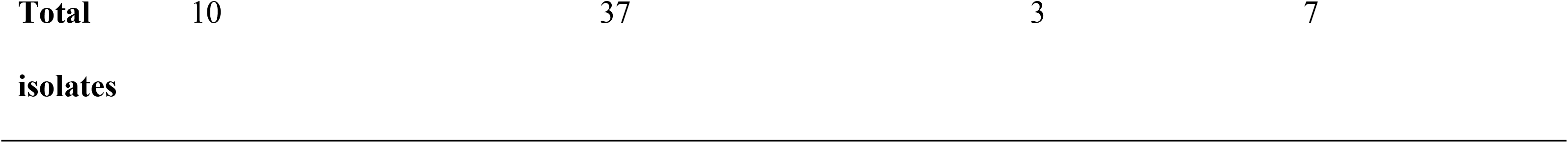
Distribution of sequence types (STs) of Group B *Streptococcus* (GBS) isolated from camel milk and extramammary body sites collected from six herds in Laikipia County, Kenya, November 2019.

**Figure 2.**
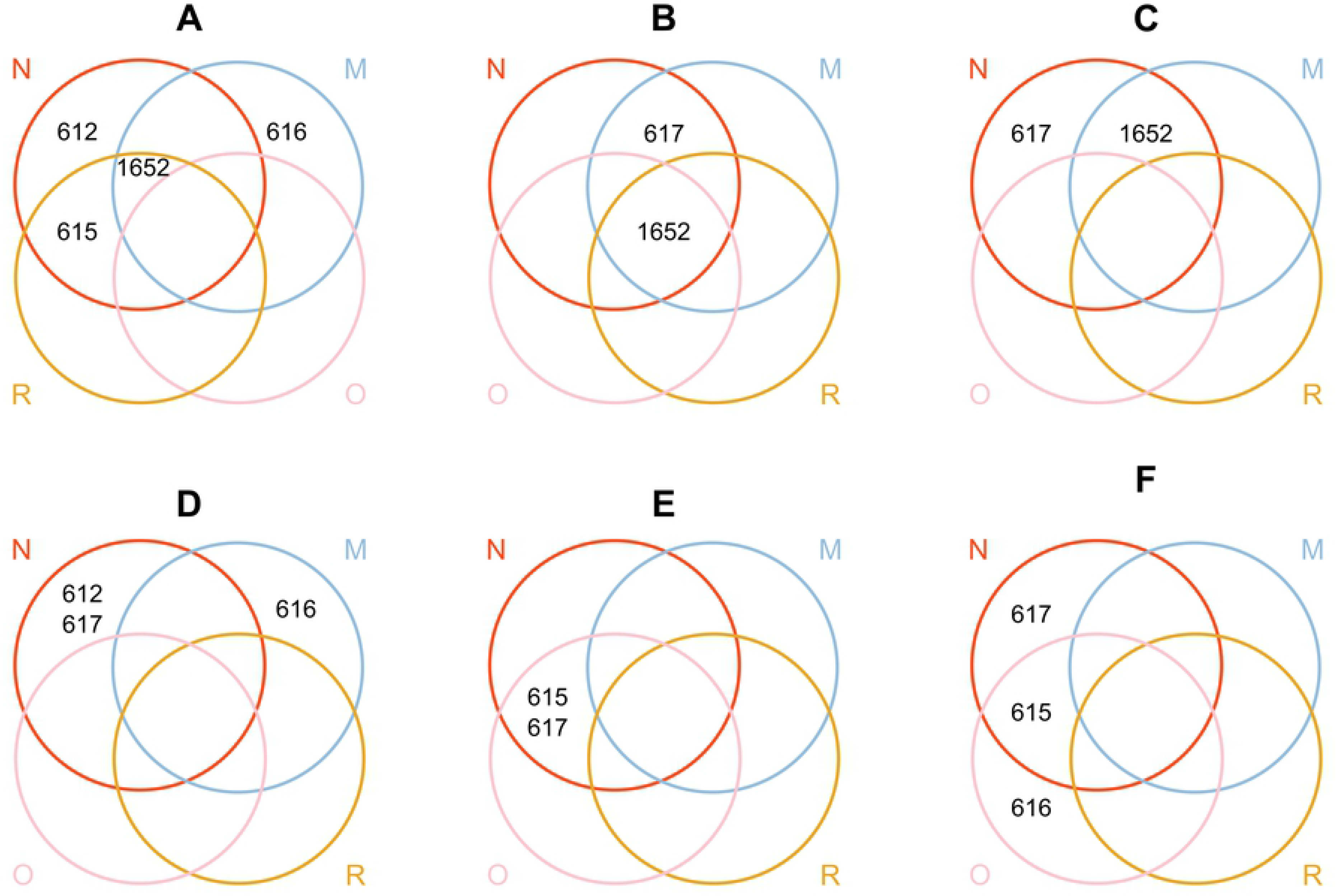
Visualisation of the herd distribution of Group B *Streptococcus* sequence types (STs) isolated from different sampling sites in six camel herds (A-F) in Laikipia County, Kenya, November 2019. Colour of the rings and capital letters correspond to sampling sites; N=nasal swab (red), R= rectal swab (yellow), O=oral swab (pink), M=milk sample (blue). Note that the position of rings for herd A is modified to allow for visualisation of the data.

Four capsular serotypes were detected *in silico*. The most common serotype was serotype VI (n=30), followed by serotype IV (n=12), serotype II (n=10) and serotype III (n=5). Serotypes II and III were only found in one ST each, whereas serotypes IV and VI were found in two STs. Except for ST617, STs were associated with a single serotype (*p*<0.001) **(Table 4)**. Serotypes matched perfectly with phylogenetic groups (**Figure 3**), with no indication of capsular switching within lineages.

**Table 4.** Distribution of molecular serotypes and sequence types (STs) among 57 Group B *Streptococcus* isolates collected from milk, nasal, rectal and oral mucosa from lactating camels and their calves in six herds in Laikipia County, Kenya, 2019.

| **Capsular serotype** | **ST612** | **ST615** | **ST616** | **ST617** | **ST1652** | **Total** |
| --- | --- | --- | --- | --- | --- | --- |
| II | - | 10 | - | - | - | 10 |
| III | - | - | 5 | - | - | 5 |
| IV | 6 | - | - | 6 | - | 12 |
| VI | - | - | - | 11 | 19 | 30 |
| **Total** | **6** | **10** | **5** | **17** | **19** | **57** |

**Figure 3.**
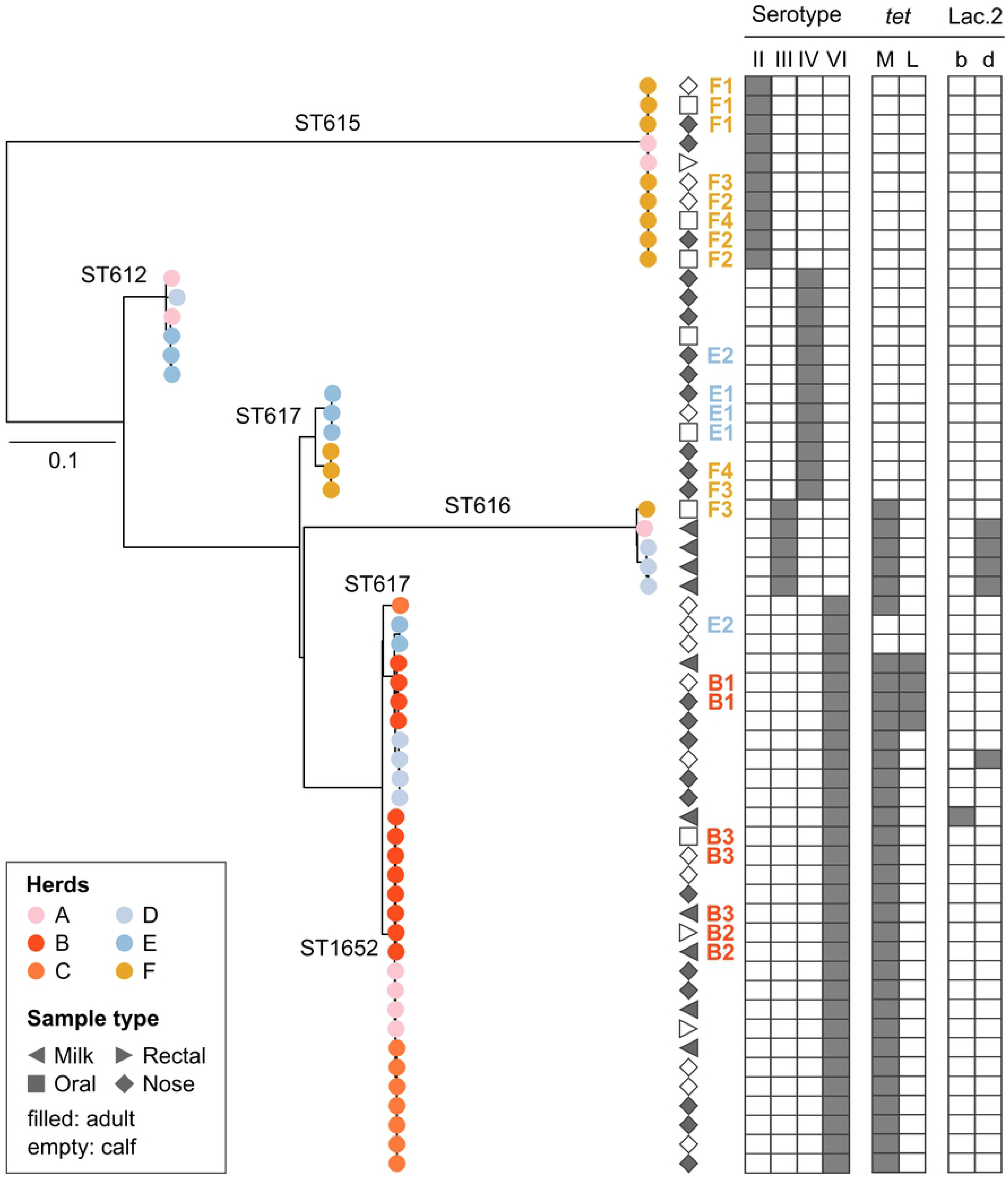
Maximum likelihood phylogenetic tree based on a core genome alignment for 57 group B *Streptococcus* isolates from milk and nasal, rectal and oral mucosa of dromedary camels in Kenya. Leaf colours indicate herd of origin with sequence types (STs) indicated on the branches. Geometric shapes correspond to sample source, while age category (adult or calf) is indicated by shading. Combinations of letters and numbers show adult-calf pairs within herds (B1, B2, B3, E1, E2, F1, F2, F3, F4) with colour corresponding to herd of origin. Grey blocks mark capsular serotype, tetracycline resistance genes, and the presence of a lactose operon in the genome (variants Lac.2b and Lac.2d). Tree was rooted at midpoint.

### Phylogenetic analysis

In the core genome phylogeny, five main lineages can be observed (**Figure 3**). Four lineages corresponded to a single ST (ST612, ST615, ST616 and ST617 respectively) and the fifth and largest lineage included ST617 and its SLV, ST1652. Two of these lineages involved isolates from only nasal and oral mucosa, while ST616 was primarily found in milk samples. To a large extent, isolates from the same herd clustered close to each other within each lineage. Nine adult-calf pairs were GBS-positive, with two or three GBS isolates per pair. For six adult-calf pairs, isolates were genetically related (B1, B2, B3, E1, F1, F2, **Figure 3**). Only two of these adult-calf pairs involved an isolate from the mother’s milk (B2 and B3). In one adult-calf pair (B2, **Figure 3**), the isolate from the milk clustered with an isolate from the rectal mucosa of the calf. In the other pair (B3), the isolate from the mother’s milk clustered with one isolate from the oral mucosa and one isolate from the nasal mucosa of the calf. The other four closely related adult-calf pairs consisted of nasal isolates from both the mother and calf. In the remaining three adult-calf pairs, isolates belonged to different lineages (E2, F3, F4, **Figure 3**).

### Antimicrobial susceptibility and lactose typing

The phenotypic susceptibility testing revealed non-WT phenotypes for tetracycline (MIC above 1 μg/mL) in 33 of 57 GBS isolates. MIC values for all other categories of antibiotics were within MIC ranges classified as WT or susceptible (for ECOFF or clinical breakpoints, respectively) with the exception of enrofloxacin, for which ECOFF and clinical breakpoint were lacking. The distribution of MIC values is shown in **Table 5.** MIC values for each isolate are presented in the **supplementary material**. The proportion of tetracycline non-WT isolates differed significantly between herds (*p*<0.001). In herd B and C, all isolates were classified as non-WT whereas in herd E, all isolates were WT. In all phenotypically tetracycline non-WT isolates, the *tet*(M) gene was found. In addition, four nasal isolates from one herd (herd D) harboured the *tet*(L) gene. No resistance genes were found in the phenotypically WT isolates. Isolates originating from milk were more likely to be tetracycline non-WT (10 out of 10) than isolates from extramammary sources (*p*=0.015). There was an association between ST and tetracycline non-WT (*p*<0.001), with all isolates belonging to ST1652 (n=19) and ST616 (n=5) being resistant.

**Table 5.** Distribution (percentage of isolates) of minimum inhibitory concentration (MIC) and prevalence of non-WT or resistance (R, shown as percentage) for Group B *Streptococcus* (n=58) isolated from lactating camels and their calves in Laikipia County, Kenya, 2019^1^.

| Test agent | % | Distribution (%) of MICs ($\mu\text{g/mL}$) | | | | | | | | | | | |
| --- | --- | --- | --- | --- | --- | --- | --- | --- | --- | --- | --- | --- | --- |
| | | ≤<br>0.03 | 0.06 | 0.125 | 0.25 | 0.5 | 1 | 2 | 4 | 8 | 16 | 32 | 64 |
| Cephalotin | 0 |  |  |  |  |  | 100 |  |  |  |  |  |  |
| Clindamycin | 0 |  |  |  |  | 100 |  |  |  |  |  |  |  |
| Enrofloxacin | NA |  |  |  |  | 70.7 | 29.3 |  |  |  |  |  |  |
| Erytromycin | 0 |  |  |  |  | 100 |  |  |  |  |  |  |  |
| Gentamicin | 0 |  |  |  |  |  |  | 5.2 | 65.5 | 29.3 |  |  |  |
| Nitrofurantoin | 0 |  |  |  |  |  |  |  |  |  | 100 |  |  |
| Penicillin | 0 | 27.6 | 72.4 |  |  |  |  |  |  |  |  |  |  |
| Tetracycline | 57 |  |  |  | 39.7 | 3.4 |  |  | 1.7 | 55.2 |  |  |  |
| Trimethoprim-Sulfamethoxazol | 0 |  |  |  | 100 |  |  |  |  |  |  |  |  |

<sup>1</sup>Unshaded cells indicate the range of concentrations tested for each antimicrobial agent. Shaded cells indicate concentrations outside the range tested for each substance. MIC equal to or lower than the lowest concentration tested for an antibiotic substance ( $\leq Y \mu\text{g/mL}$ ), is given as a percentage at the lowest tested concentration. Blank unshaded cells indicate lack of isolates with that MIC. Bold vertical lines indicate epidemiological cut-off values retrieved from the European Committee on Antimicrobial Susceptibility Testing or, for cephalotin and trimethoprim-sulfamethoxazol, clinical breakpoints from SVA.

Only six isolates from three herds (A, B and D) were lactose-positive based on phenotyping, PCR, and genomic analysis, with full agreement between methods. Lactose fermentation was observed at the first visual assessment after 24 hours of incubation. Five lactose fermenting isolates originated from milk, and one from the nasal mucosa of a calf. Four lactose fermenting isolates belonged to ST616, and one each to ST617 and ST1652. There was an association between lactose fermentation and ST, with ST616 being overrepresented among lactose fermenting strains (4 out of 6) (*p*=0.05), but no association between lactose operon variant (Lac.2b, n=1; Lac2.d, n=5) and sample type.

## Discussion

### Nasal carriage of GBS is highly prevalent in camels

We found a high prevalence of GBS on the nasal mucosa of apparently healthy animals, similar to the findings by Younan and Bornstein [12], but in contrast to the low prevalence described in neighbouring counties [38]. Furthermore, isolates from the nasal mucosa showed a high level of genetic diversity, with isolates belonging to all STs identified in the study, with the exception of ST616, an ST which is significantly associated with the mammary gland [39]. These findings suggest that nasal colonisation of GBS in healthy camels is common, in contrast to the situation in humans and cattle. In humans, prevalence of nasopharyngeal carriage is estimated at around 10% in different geographic regions [29,40,41]. The observation of a camel-calf pair with a shared nasal GBS type suggest nose to nose contact may be a route of transmission. Reports of GBS in respiratory tract infections in camels suggest that the nasal cavity may serve as a reservoir for opportunistic GBS infections [12,42].

In humans, rectal colonisation is the most common form of GBS carriage [29,43] and widely recognized as a source of sporadic disease in infants [44]. By contrast, we observed a very low isolation rate from the rectal mucosa in camels, similar to a previous report where none of 25 camel rectal samples tested positive for GBS [12], and to the situation in cattle [9,10]. The finding of two genetically closely related isolates, originating from a rectal sample from a suckling calf and milk from its mother, suggests the probable passage of GBS through the gastrointestinal tract in camels. It is, however, not possible to say whether these findings are the results of ingestion of GBS infected milk by the calf or if it is indicative of rectal colonisation. Vaginal colonisation was not observed in our study, suggesting absence or a low prevalence, compatible with a reported prevalence of vaginal GBS carriage in camels of 1 to 21% [12]. Vaginal colonisation is also relatively rare in cattle [10]. By contrast, and mirroring the prevalence of rectal colonisation, vaginal prevalence of GBS in humans is much higher, generally estimated to be in the range of 20 to 40% [44]. Thus, our data imply that the niche-predilection of GBS differs between host species, with preferred sites being the rectum and urogenital tract in humans, the mammary gland in cattle, and the nasal mucosa in camels. For *Staphylococcus aureus*, another important mastitis pathogen of ruminants and camelids, similar differences in niche-predilection have been described, with the mammary gland being the most common site of *S. aureus* isolation in cattle, and the nose the most common site in sheep [45]. Two STs; ST617 and its SLV, ST1652, were isolated from all positive sampling sites, demonstrating that this niche-predilection is not exclusive. In a previous study, ST617 was isolated predominantly from wound infections or abscesses but also from cases of mastitis [15], suggesting that skin, soft tissue and intramammary infections, like respiratory infections, may be opportunistic. Another striking difference between camel isolates and those from humans and cattle is the distribution of serotypes. Serotype VI was the most common serotype in the current study, but this serotype is very rare among human GBS, to the point where it is not included in current human GBS vaccine trials [46]. Serotype IV was the second most common type in camels in our study. This is an emerging serotype in humans, i.e. historically of limited relevance but increasingly recognised as a cause of disease [47]. Some STs with serotype IV are found in humans as well as cattle, notably GBS ST196 [48], but there is no evidence for occurrence of any of the camel-associated STs with serotype IV in humans. Strikingly, serotypes Ia, Ib and V, which are common in human and bovine GBS [48,49] were absent from the camel GBS population in our study.

### Extramammary GBS strains are a potential source of mastitis in camels

Only four of ten GBS-positive milk samples belonged to ST616, which has previously been associated with mastitis [39]. The remaining milk isolates belonged to STs that were predominantly found in extramammary sources. Thus, GBS mastitis in camels may have a mixed epidemiology, comprising transmission from udder to udder (“contagious transmission”) and transmission from extramammary sources (“environmental transmission”). Although GBS was isolated from rectal samples on a few occasions in this study, this is not a likely route for within-herd transmission in camel herds, considering that camels are mostly kept in an arid environment and that camel faeces are unlikely to contaminate the environment due to their dry and firm consistence, in contrast to the situation for high-producing or housed dairy cattle. Transmission from skin or mucosa, as described for *S. aureus* in cattle [50,51], seems more likely considering our findings. The existence of extramammary GBS on the nasal and oral mucosa of calves also points towards calves acting as reservoirs for GBS, which could contribute to transmission of extramammary isolates to the udder, as shown for *S. aureus* in sheep [45]. In contrast to what has previously been reported [52], we observed cross-suckling on a few occasions. It has been suggested that suckling calves can cause contamination of the camel udder [14] and contribute to udder-to-udder transmission of GBS.

Herds E and F had comparatively high isolation rates of GBS from the nasal and oral mucosa of calves but no GBS was found in milk. This observation may result from our study design, whereby not all camels’ udders were sampled, but only those with CMT ≥ 2 in at least one quarter. GBS has previously been isolated from quarters without an elevated cell count [7] and similarly, from dairy cows with a composite somatic cell count of <200,000 cells/mL [53]. Hence, it is possible that camels with GBS mastitis could have passed undetected in this study. It may explain how ST616 could be detected on the oral mucosa of a calf without having been detected in the milk of its mother (pair F3, **Figure 3**). This limitation in animals selected for milk sampling was unfortunately necessary to ensure animal owner compliance with the sampling procedure.

The association between specific GBS types and intramammary infection was thought to be due to the presence of the lactose operon (Lac.2), both in cattle [45] and in camels [39]. In a study on camels from a different region in Kenya and with a different production system, more than 80% of GBS isolates from milk harboured Lac.2 [39]. In the current study, Lac.2 was only detected in 40% of milk isolates. The ability to ferment lactose likely offers GBS an evolutionary advantage in adaption to the mammary gland, however, it appears that for camel GBS this is not a necessary feature to establish intramammary infections. As this study was cross-sectional, we cannot determine whether the presence of a Lac.2 operon was associated with chronicity of infection.

### The local context affects the epidemiology of GBS in camel herds

The phylogenetic analysis revealed distinct herd clusters, reflecting how ranches and smallholder herds are relatively closed management systems. This outcome is quite distinct from previous results for milk isolates from traditional pastoralist herds, where highly similar strains were identified in multiple herds [39]. Even on ranches and in smallholder systems, however, between-herd transmission may be possible, as illustrated by the findings of genetically related isolates in herd A, E and F. This can likely be explained by known epidemiological risk factors for between-herd transmission, such as the owners of two of these herds being close neighbours and having frequent contact, and household members from herd F being employed at the ranch of herd A.

Prevalence of tetracycline non-WT among GBS isolates was herd-associated and ST-associated, with non-WT being present in all isolates from two herds compared to none or almost none from two other herds. The *tet*(M) gene, one of the most common genes for tetracycline resistance, was found in all non-WT isolates. This gene, which is carried by integrative conjugative elements, has been described in GBS in camels as well as other host species, notably humans [15,47]. In four isolates, we detected the *tet*(L) gene which has not previously been reported in camel GBS. The observed differences in susceptibility patterns among isolates originating from different herds herds and STs are likely due to the combination of selective pressure from antimicrobial use, horizontal gene transfer of resistance genes between GBS lineages, and transmission of GBS lineages with or without resistance genes within and between herds. Antibiotic sales and use in Kenya are largely unmonitored [54], but antimicrobials, particularly tetracycline, are reportedly a common treatment option against infections in camels [55,56]. Interestingly, isolates originating from milk were more likely to be tetracycline resistant than isolates from other sampling sites. This is in agreement with previous findings of high levels of tetracycline resistance in GBS from milk [7] and tetracycline resistance being less common in GBS from the nasal mucosa in camels [38]. In cattle, treatment of mastitis with tetracycline has been shown to be inefficient and results in subtherapeutic concentrations in the udder, something that would promote development or acquisition of resistance in udder pathogens [57,58]. For example, exposure to tetracycline and other subtherapeutic concentrations of ribosome-targeting antibiotics, e.g. macrolides, lincosamides, and streptogramins may induce transfer of Tn*916* [59], a mobile genetic element that facilitates horizontal transfer of resistance in GBS from humans and camels [15,49].

## Conclusions

In conclusion, the prevalence of GBS in the nares of camels was higher than observed in any other GBS host species. In addition, occasional oral carriage and faecal shedding were detected but no vaginal carriage, in stark contrast to the situation in humans, were rectal and vaginal carriage are far more common than oropharyngeal colonisation, or the situation in cattle, where all extramammary detection of GBS is relatively rare. The serotype distribution of GBS was markedly different between camels and its other major mammalian hosts (humans and cattle), adding further evidence that the ecology of GBS in camels is unique. Of five STs that were detected, only ST616 was significantly associated with milk, but the majority of isolates from milk belonged to STs that were also detected in extramammary sources, suggesting that they may play an important role in the epidemiology of mastitis in camels, which is in contrast to the situation in cattle.

## Acknowledgements

The authors would like to extend their gratitude to all pastoralists and camel ranch owners that participated in this study. We thank Mpala Research Centre for assistance in organising the field work. We also thank Mattias Myrenås and Oskar Nilsson (Department of Animal Health and Antimicrobial Strategies, SVA, Sweden) for help with the PCR-analysis and for guidance on antimicrobial susceptibility testing, respectively.

## Supporting information

**S1. Dataset. Authors’ original data for 57 GBS-isolates collected from camels in Laikipia County, Kenya, October - November, 2019**. The dataset includes accession numbers available at European Nucleotide Archive (ENA), information regarding herd identity, source of the isolate, sequence type (ST), capsular serotype, minimum inhibitory concentration (MIC) values for phenotypic antimicrobial resistance testing, presence/absence of antimicrobial resistance (AMR) genes, phenotypic lactose fermentation results, Lac.2 PCR results, presence/absence of the Lac.2 operon genotypes and herd-coordinates.

## References

1. Catley A, Lind J, Scoones I. The futures of pastoralism in the Horn of Africa: pathways of growth and change. Rev Sci Tech OIE. 2016 Aug 1;35(2):389–403.

2. Bekele T, Lundeheim N, Dahlborn K. Milk production and feeding behavior in the camel (*Camelus dromedarius*) during 4 watering regimens. J Dairy Sci. 2011 Mar 1;94(3):1310–7.

3. Faye B, Chaibou M, Gilles V. Integrated impact of climate change and socioeconomic development on the evolution of camel farming systems. Br J Environ Clim Change. 2012 Oct 6;2:227–44.

4. Kagunyu AW, Wanjohi J. Camel rearing replacing cattle production among the Borana community in Isiolo County of Northern Kenya, as climate variability bites. Pastoralism. 2014 Aug 28;4(1):13.

5. Anderson DM, Elliott H, Kochore HH, Lochery E. Camel herders, middlewomen, and urban milk bars: the commodification of camel milk in Kenya. J East Afr Stud. 2012 Aug 1;6(3):383–404.

6. Farah Z, Mollet M, Younan M, Dahir R. Camel dairy in Somalia: Limiting factors and development potential. Livest Sci. 2007 Jun 1;110(1):187–91.

7. Seligsohn D, Nyman A-K, Younan M, Sake W, Persson Y, Bornstein S, et al. Subclinical mastitis in pastoralist dairy camel herds in Isiolo, Kenya: Prevalence, risk factors, and antimicrobial susceptibility. J Dairy Sci. 2020 May 1;103(5):4717–31.

8. Matofari JW, Younan M, Mwatha EW, Okemo PO. Microorganisms associated with sub-clinical mastitis in the Kenyan camel (C*amelus dromedarius*). J Trop Microbiol Biotechnol. 2003;2(1):11–6.

9. Cobo-Ángel C, Jaramillo-Jaramillo AS, Lasso-Rojas LM, Aguilar-Marin SB, Sanchez J, Rodriguez-Lecompte JC, et al. *Streptococcus agalactiae* is not always an obligate intramammary pathogen: Molecular epidemiology of GBS from milk, feces and environment in Colombian dairy herds. PLOS ONE. 2018 Dec 10;13(12):e0208990.

10. Jørgensen HJ, Nordstoga AB, Sviland S, Zadoks RN, Sølverød L, Kvitle B, et al. *Streptococcus agalactiae* in the environment of bovine dairy herds – rewriting the textbooks? Vet Microbiol. 2016 Feb 29;184:64–72.

11. Manning SD, Springman AC, Million AD, Milton NR, McNamara SE, Somsel PA, et al. Association of group B *Streptococcus* colonization and bovine exposure: a prospective cross-sectional cohort study. PLOS ONE. 2010 Jan 20;5(1):e8795.

12. Younan M, Bornstein S. Lancefield group B and C streptococci in East African camels (*Camelus dromedarius*). Vet Rec. 2007 Mar 10;160(10):330–5.

13. Schalm OW. *Streptococcus agalactiae* in the udders of heifers at parturition traced to suckle among calves. Cornell Vet. 1942;32:49–60.

14. Noor IM, Guliye AY, Tariq M, Bebe BO. Assessment of camel and camel milk marketing practices in an emerging peri-urban production system in Isiolo County, Kenya. Pastor Res Policy Pract. 2013 Dec 2;3(1):28.

15. Fischer A, Liljander A, Kaspar H, Muriuki C, Fuxelius H-H, Bongcam-Rudloff E, et al. Camel *Streptococcus agalactiae* populations are associated with specific disease complexes and acquired the tetracycline resistance gene *tet*M via a Tn*916*-like element. Vet Res. 2013 Oct 1;44(1):86.

16. Georgiadis NJ, Olwero JGN, Ojwang’ G, Romañach SS. Savanna herbivore dynamics in a livestock-dominated landscape: I. Dependence on land use, rainfall, density, and time. Biol Conserv. 2007 Jul 1;137(3):461–72.

17. Kenya National Bureau of Statistics & County Government of Laikipia. County Statistical Abstract - Laikipia County. Kenya National Bureau of Statistics; 2019 [cited 2021 Mar 9]. Available from: https://www.laikipia.go.ke/assets/file/3cc125a1-laikipia-county-statistical-abstract.pdf

18. Volpato G, King EG. From cattle to camels: trajectories of livelihood adaptation and social-ecological resilience in a Kenyan pastoralist community. Reg Environ Change. 2019 Mar 1;19(3):849–65.

19. Tinggren S. Udder health inflammatory markers in camel milk (*Camelus dromedarius*) and milk yield [Internet]. Uppsala: SLU, Dept. of Clinical Sciences; 2019 [cited 2021 Feb 24]. Available from: https://stud.epsilon.slu.se/14383/

20. Schalm OW, Noorlander DO. Experiments and observations leading to development of the California mastitis test. J Am Vet Med Assoc. 1957 Mar 1;130(5):199–204.

21. Klastrup O, Madsen PS. Nordiske rekommendationer vedrorende mastitisundersogelser af kirtelprover (Nordic recommendations concerning mastitis control of quarter samples). Nord Veterinaermed. 1974;26:197–204.

22. Christie K, Atkins NE, Munch-Petersen E. A note on a lytic phenomenon shown by group B streptococci. Aust J Exp Biol Med Sci. 1944;22(3):197–200.

23. Bizzini A, Durussel C, Bille J, Greub G, Prod’hom G. Performance of matrix-assisted laser desorption ionization-time of flight mass spectrometry for identification of bacterial strains routinely isolated in a clinical microbiology laboratory. J Clin Microbiol. 2010 Jan 5;48(5):1549–54.

24. Clinical and Laboratory Standards Institute (CLSI). Performance standards for antimicrobial susceptibility testing: M100, S27. Vol. 28. Wayne, PA; 2017.

25. Gupta SK, Padmanabhan BR, Diene SM, Lopez-Rojas R, Kempf M, Landraud L, et al. ARG-ANNOT, a new bioinformatic tool to discover antibiotic resistance genes in bacterial genomes. Antimicrob Agents Chemother. 2014 Jan 1;58(1):212–20.

26. Richards VP, Lang P, Pavinski Bitar PD, Lefébure T, Schukken YH, Zadoks RN, et al. Comparative genomics and the role of lateral gene transfer in the evolution of bovine adapted *Streptococcus agalactiae*. Infect Genet Evol. 2011 Aug 1;11(6):1263–75.

27. Camacho C, Coulouris G, Avagyan V, Ma N, Papadopoulos J, Bealer K, et al. BLAST+: architecture and applications. BMC Bioinformatics. 2009 Dec 15;10(1):421.

28. Crestani C. The fall and rise of group B *Streptococcus* in dairy cattle: reintroduction due to human-to-cattle host jumps? BioRxiv [Preprint]. 2021. bioRxiv:440740.440740v1.[posted 2021 April 22; cited 2021 April 23]. doi:10.1101/2021.04.21.440740

29. Sørensen UBS, Klaas IC, Boes J, Farre M. The distribution of clones of *Streptococcus agalactiae* (group B streptococci) among herdspersons and dairy cows demonstrates lack of host specificity for some lineages. Vet Microbiol. 2019 Aug 1;235:71–9.

30. Seemann T. Prokka: rapid prokaryotic genome annotation. Bioinforma Oxf Engl. 2014 Jul 15;30(14):2068–9.

31. Smeds L, Künstner A. ConDeTri - a content dependent read trimmer for Illumina data. PLOS ONE. 2011 Oct 19;6(10):e26314.

32. Bankevich A, Nurk S, Antipov D, Gurevich AA, Dvorkin M, Kulikov AS, et al. SPAdes: a new genome assembly algorithm and its applications to single-cell sequencing. J Comput Biol. 2012 Apr 16;19(5):455–77.

33. Gurevich A, Saveliev V, Vyahhi N, Tesler G. QUAST: quality assessment tool for genome assemblies. Bioinformatics. 2013 Apr 15;29(8):1072–5.

34. Cineros JLB, Lund O. KmerFinderJS: A client-server method for fast species typing of bacteria over slow Internet connections. bioRxiv. 2017 Jun 2;145284.

35. Inouye M, Dashnow H, Raven L-A, Schultz MB, Pope BJ, Tomita T, et al. SRST2: Rapid genomic surveillance for public health and hospital microbiology labs. Genome Med. 2014 Nov 20;6(11):90.

36. Metcalf BJ, Chochua S, Gertz RE, Hawkins PA, Ricaldi J, Li Z, et al. Short-read whole genome sequencing for determination of antimicrobial resistance mechanisms and capsular serotypes of current invasive *Streptococcus agalactiae* recovered in the USA. Clin Microbiol Infect. 2017 Aug 1;23(8):574.e7–574.e14.

37. Kozlov AM, Darriba D, Flouri T, Morel B, Stamatakis A. RAxML-NG: a fast, scalable and user-friendly tool for maximum likelihood phylogenetic inference. Bioinformatics. 2019 Nov 1;35(21):4453–5.

38. Mutua JM, Gitao CG, Bebora LC, Mutua FK. Antimicrobial resistance profiles of bacteria isolated from the nasal cavity of camels in Samburu, Nakuru, and Isiolo Counties of Kenya. J Vet Med. 2017;2017:1216283.

39. Seligsohn D, Crestani C, Forde TL, Chenais E, Zadoks RN. Genomic analysis of group B *Streptococcus* from milk demonstrates the need for improved biosecurity: a cross-sectional study of pastoralist camels in Kenya. BMC Microbiology, forthcoming.

40. Cobo-Àngel CG, Jaramillo-Jaramillo AS, Palacio-Aguilera M, Jurado-Vargas L, Calvo-Villegas EA, Ospina-Loaiza DA, et al. Potential group B *Streptococcus* interspecies transmission between cattle and people in Colombian dairy farms. Sci Rep. 2019 Dec;9(1):14025.

41. Foster-Nyarko E, Kwambana B, Aderonke O, Ceesay F, Jarju S, Bojang A, et al. Associations between nasopharyngeal carriage of group B *Streptococcus* and other respiratory pathogens during early infancy. BMC Microbiol. 2016 May 27;16(1):97.

42. Motlová J, Straková L, Urbásková P, Sak P, Sever T. Vaginal & rectal carriage of *Streptococcus agalactiae* in the Czech Republic: Incidence, serotypes distribution & susceptibililty to antibiotics. Indian J Med Res. 2004 Jun 1;119 Suppl:84–7.

43. Wernery U, Kinne J, Anas S, John J. Adhesive pleurisy of both lungs in a dromedary camel caused by *Streptococcus agalactiae*: A case report. J Camel Pract Res. 2018;25(3):311.

44. Le Doare K, Heath PT. An overview of global GBS epidemiology. Vaccine. 2013 Aug 28;31:D7–12.

45. Mørk T, Kvitle B, Jørgensen HJ. Reservoirs of *Staphylococcus aureus* in meat sheep and dairy cattle. Vet Microbiol. 2012 Feb 24;155(1):81–7.

46. Lyhs U, Kulkas L, Katholm J, Waller KP, Saha K, Tomusk RJ, et al. *Streptococcus agalactiae* serotype IV in humans and cattle, Northern Europe. Emerg Infect Dis. 2016 Dec;22(12):2097–103.

47. Richards VP, Velsko IM, Alam MT, Zadoks RN, Manning SD, Pavinski Bitar PD, et al. Population gene introgression and high genome plasticity for the zoonotic pathogen *Streptococcus agalactiae*. Mol Biol Evol. 2019 Nov 1;36(11):2572–90.

48. Absalon J, Segall N, Block SL, Center KJ, Scully IL, Giardina PC, et al. Safety and immunogenicity of a novel hexavalent group B *Streptococcus* conjugate vaccine in healthy, non-pregnant adults: a phase 1/2, randomised, placebo-controlled, observer-blinded, dose-escalation trial. Lancet Infect Dis. 2021 Feb 1;21(2):263–74.

49. Teatero S, Athey TBT, Caeseele PV, Horsman G, Alexander DC, Melano RG, et al. Emergence of serotype IV group B *Streptococcus* adult invasive disease in Manitoba and Saskatchewan, Canada, is driven by clonal sequence type 459 strains. J Clin Microbiol. 2015 Sep 1;53(9):2919–26.

50. Capurro A, Aspán A, Ericsson Unnerstad H, Persson Waller K, Artursson K. Identification of potential sources of *Staphylococcus aureus* in herds with mastitis problems. J Dairy Sci. 2010 Jan 1;93(1):180–91.

51. Zadoks RN, Leeuwen WB van, Kreft D, Fox LK, Barkema HW, Schukken YH, et al. Comparison of *Staphylococcus aureus* isolates from bovine and human skin, milking equipment, and bovine milk by phage typing, pulsed-field gel electrophoresis, and binary typing. J Clin Microbiol. 2002 Nov 1;40(11):3894–902.

52. Packer C, Lewis S, Pusey A. A comparative analysis of non-offspring nursing. Anim Behav. 1992 Feb 1;43(2):265–81.

53. Mahmmod YS, Klaas IC, Katholm J, Lutton M, Zadoks RN. Molecular epidemiology and strain-specific characteristics of *Streptococcus agalactiae* at the herd and cow level. J Dairy Sci. 2015 Oct;98(10):6913–24.

54. Heffernan C, Misturelli F. The delivery of veterinary services to the poor: findings from Kenya [Internet]. Reading, UK: Veterinary and Economics Research Unit, Department of Agriculture, University of Reading.; 2000 [cited 2020 Dec 18]. Available from: https://agris.fao.org/agris-search/search.do?recordID=GB2013202352

55. Lamuka PO, Njeruh FM, Gitao GC, Abey KA. Camel health management and pastoralists’ knowledge and information on zoonoses and food safety risks in Isiolo County, Kenya. Pastoralism. 2017 Aug 2;7(1):20.

56. Younan M. Traitement parentéral de mammites à *Streptococcus agalactiae* chez le dromadaire (*Camelus dromedarius*) au Kenya. Rev D’élevage Médecine Vét Pays Trop. 2002 Mar 1;55(3):177–81.

57. Gruet P, Maincent P, Berthelot X, Kaltsatos V. Bovine mastitis and intramammary drug delivery: review and perspectives. Adv Drug Deliv Rev. 2001;50(3):245–59.

58. Lents CA, Wettemann RP, Paape MJ, Vizcarra JA, Looper ML, Buchanan DS, et al. Efficacy of intramuscular treatment of beef cows with oxytetracycline to reduce mastitis and to increase calf growth. J Anim Sci. 2002 Jun;80(6):1405–12.

59. Scornec H, Bellanger X, Guilloteau H, Groshenry G, Merlin C. Inducibility of Tn*916* conjugative transfer in *Enterococcus faecalis* by subinhibitory concentrations of ribosome-targeting antibiotics. J Antimicrob Chemother. 2017 Oct 1;72(10):2722–8.

